# The convergence and divergence of episodic and semantic functions across lateral parietal cortex

**DOI:** 10.1101/2020.12.08.416123

**Authors:** Gina F. Humphreys, JeYoung Jung, Matthew A. Lambon Ralph

**Author notes:** <u>Address for correspondence</u>: Dr. Gina Humphreys or Prof. Matt Lambon Ralph, MRC Cognition and Brain Sciences Unit, University of Cambridge, 15 Chaucer Road, Cambridge, CB2 7EF, UK.

## Abstract

Several decades of neuropsychological and neuroimaging research have highlighted the importance of lateral parietal cortex (LPC) across a myriad of cognitive domains. Yet, despite the prominence of this region the underlying function of LPC remains unclear. Two domains that have placed particular emphasis on LPC involvement are semantic memory and episodic memory retrieval. From each domain, sophisticated models have been proposed as to the underlying function, as well as the more domain-general assumption that LPC is engaged by any form of internally-directed cognition (episodic and semantic retrieval both being examples if this process). Here we directly address these alternatives using a combination of fMRI, functional connectivity and DTI white-matter connectivity data. The results show that ventral LPC (angular gyrus) was positively engaged during episodic retrieval but disengaged during semantic memory retrieval. In addition, the level of activity negatively varied with task difficulty in the semantic task whereas episodic activation was independent of difficulty. In contrast, dorsal LPC (intraparietal sulcus) showed domain general activation that was positively correlated with task difficulty. In terms of functional and structural connectivity, a dorsal-ventral and anterior-posterior gradient of connectivity was found to different processing networks (e.g., mid-angular gyrus (AG) connected with episodic retrieval). We propose a unifying model in which LPC as a whole might share a common underlying neurocomputation (e.g., multimodal buffering) with variations in the emergent, expressed cognitive functions across subregions arising from differences in the underlying white matter connectivity.

## Introduction

Several decades of neuropsychological and neuroimaging research have highlighted the importance of lateral parietal cortex (LPC) across a myriad of cognitive domains (Corbetta and Shulman 2002; Wagner et al. 2005; Binder et al. 2009; Cabeza et al. 2012; Humphreys and Lambon Ralph 2015; Sestieri et al. 2017). It also forms a core part of the default mode network (DMN), a network that often deactivates during task-performance (Raichle et al. 2001; Buckner et al. 2008; Humphreys et al. 2015) Yet, despite the prominence of this region in basic and clinical research, the underlying function of LPC remains unclear. Part of this confusion may reflect that cognitive neuroscience research is often focused and organised by the cognitive domain of interest. As a result, multiple cognitive domains have been associated with the LPC with minimal cross-talk between these separate literatures and resultant cognitive neuroscience theories (with a few notable exceptions: (Corbetta and Shulman 2002; Cabeza *et al*. 2012; Humphreys and Lambon Ralph 2015; Rugg and King 2018; Renoult et al. 2019). Whilst these theories are often sophisticated and are based on a wealth of domain-specific data, they fail to explain both the wide variety of functions that appear to recruit this region and, thus, what the core underpinning neurocomputations might be. There are two types of answer to this fundamental question (Humphreys and Lambon Ralph 2015; Humphreys, Lambon Ralph, et al. 2020) 1) a form of ‘neuromarquetry’ in which LPC contains numerous sub-regions each recruited by different tasks and serve distinct underlying cognitive functions; or 2) multiple cognitive activities rely, in common, upon a small number of underlying computations that arise from the LPC and its patterns of connectivity to the wider neural network (Caspers et al. 2008; Uddin et al. 2010; Caspers et al. 2011; Cloutman et al. 2013). Given that the LPC is an anatomically heterogeneous region in terms of cytoarchitecture and functional/structural connectivity across LPC (Caspers *et al*. 2008; Uddin *et al*. 2010; Caspers *et al*. 2011; Mars et al. 2011; Cloutman *et al*. 2013), at least some variability in cognitive function across LPC might be expected. Fathoming the nature of LPC neurocomputations (and by extension other higher cortical regions) will require a sophisticated approach. Rather than focus solely upon an individual cognitive domain, it will be necessary to 1) combine data across multiple higher cortical functions, and 2) consider how varying functional input might modulate the activation pattern across tasks.

The current study therefore had two primary aims. First, to determine the underlying LPC function by directly comparing two cognitive domains (semantic and episodic memory) that are traditionally associated with the LPC, as well as the more domain-general hypothesis that LPC is engaged by any form of internally-directed cognition (episodic and semantic retrieval both being examples of this process). Despite being associated with the LPC (and, in particular, the angular gyrus) throughout long neuropsychological and functional neuroimaging literatures, these domains have rarely been directly compared in the same group of participants (and those that have been conducted have suffered from some important limitations). The second aim was to determine the extent to which variations in the expressed cognitive functions across LPC subregions directly reflect their varying input. This was achieved by directly mapping the correspondence between task fMRI data, and both functional and DTI white-matter connectivity measures.

### Domain-specific theories

Many LPC theories and datasets have focussed on individual cognitive domains. The semantic hypothesis has a very long history and tradition in both neuropsychology and from the start of functional neuroimaging. According to this proposal, the ventral LPC, specifically the angular gyrus (AG), acts as a semantic-hub which stores multi-modal semantic information (Geschwind 1972; Binder *et al*. 2009), in a similar manner to the anterior temporal lobe (ATL) (Lambon Ralph et al. 2017). The observations of concrete>abstract and words>nonword differences in the AG found in individual studies and in formal meta-analyses form a cornerstone of evidence for this proposal. In contrast, numerous studies have also observed AG responses during episodic retrieval, with activation typically correlating with the vividness of the memory retrieved. This has led to the suggestion that LPC might act as some form of episodic buffer (Wagner *et al*. 2005; Vilberg and Rugg 2008). The fact that these two very different components of human long-term memory seem to involve the same brain LPC region has received little attention despite the very large and growing number of studies on each one. In fact, the situation for episodic and semantic memory is a worked example of the broader challenge – namely that many different cognitive domains have been implicated and do overlap in the LPC, yet only a few explanations have been offered for this multi-domain maelstrom (Humphreys and Lambon Ralph 2015; Rugg and King 2018; Renoult *et al*. 2019; Humphreys, Lambon Ralph, *et al*. 2020)).

### Domain-general theories

A handful of research groups have noted this confluence of multiple cognitive functions in the LPC. The resultant theories argue that some LPC processes might be domain-general in nature and represent more fundamental neurocomputations that are required by multiple cognitive activities (Corbetta and Shulman 2002; Walsh 2003; Cabeza *et al*. 2012; Fedorenko et al. 2013; Humphreys and Lambon Ralph 2015, 2017). For instance, there is good evidence to suggest that a common region within dorsal LPC (intraparietal sulcus (IPS)) forms part of a “multiple demand network” and is recruited as part of a fronto-parietal network for executive processing (Fedorenko *et al*. 2013; Humphreys and Lambon Ralph 2015, 2017). Going further, several theories suggest that dorsal LPC (dLPC) within the IPS, and ventral LPC (vLPC) serve counterpointed functions that are utilised across cognitive domains. The dLPC has been implicated in any task that requires top-down attentional control whereas ventral areas are automatically recruited for bottom-up attention (Corbetta and Shulman 2002) Relatedly, dLPC is involved in tasks that require externally-directed attention, whereas vLPC is active during internally-directed attention, as is the case when retrieving semantic and/or episodic memories, as well as other processes such as future planning or self-projection (Buckner *et al*. 2008; Andrews-Hanna 2012). Since these internally-directed processes are not required during most fMRI tasks the vLPC is deactivated, hence its involvement in the DMN.

### Parietal Unified Connectivity-biased Computation (PUCC) model

The Parietal Unified Connectivity-biased Computation (PUCC) model takes a cross-domain perspective of LPC function. A central underlying idea is that the expressed cognitive function of each region will reflect the product of its local neurocomputation and its input/output connections (because the connections constrain what forms of information the neurocomputation acts upon). Thus PUCC is based on two key assumptions. First, the local neurocomputation is considered to be constant across the wider LPC, and provides the basis for online, multi-sensory buffering across modalities (as well as multi-modal combinations) of input. A multi-modal convergent buffer is important for bringing together multiple inputs in order to process time-extended behaviours, such as remembering an episodic event, narrative speech comprehension or sequential object use (Geschwind 1965; Damasio 1989; Botvinick and Plaut 2004, 2006). A number of prominent parallel distributed processing (PDP) computational models have shown that the addition of recurrent feedback loops allows a model to ‘buffer’ verbal or nonverbal spatiotemporal input (Elman nets: (Elman 1990)) in support of time-extended verbal and nonverbal behaviours (McClelland et al. 1989; Botvinick and Plaut 2004; Ueno et al. 2011). A “buffering-type” function is consistent, and indeed part inspired by more domain-specific buffer models of LPC function (Baddeley 2000; Wagner *et al*. 2005; Vilberg and Rugg 2008), as well as a “working-memory” type system in dorsal LPC (Pessoa et al. 2002; Humphreys and Lambon Ralph 2015).

The second key assumption of PUCC is that whilst the local neurocomputation may be constant across LPC, the ‘expressed’ task contribution of each LPC subregion will be influenced by its long-range connections. Thus, even on an assumption that the local buffering computation might be the same throughout the LPC, the types and forms of information being buffered will reflect the inputs and outputs to each subregion. This tenet is observed in various implemented computational models which have shown that the involvement of a processing unit to each cognitive activity is moulded both by its local computation, as well its connectivity to different input/output information sources (‘connectivity-constrained cognition – C3’: [28-30]). In terms of underlying architecture, anatomical evidence suggests that there are variations in cytoarchitecture and functional/structural connectivity across LPC (Caspers *et al*. 2008; Uddin *et al*. 2010; Caspers *et al*. 2011; Cloutman *et al*. 2013). For instance, the dorsal LPC is known to connect with the frontal executive network, whereas ventral LPC connects with a distributed set of regions associated with the DMN, saliency network, language network etc. (Vincent et al. 2008; Spreng et al. 2010; Uddin *et al*. 2010; Power et al. 2011; Lee et al. 2012; Cloutman *et al*. 2013; Power and Petersen 2013; Yeo et al. 2013). Consequently, at least some variability in cognitive function across LPC might be expected.

Therefore, according to PUCC, whilst LPC as a whole might share a common underlying neurocomputation (e.g., multimodal buffering), variations in task-activation across subregions will arise due to differences in the underlying input (e.g. visual, verbal, spatial, executive etc.) (Humphreys and Lambon Ralph 2015). Indeed, we have previously shown that the profile of functional activation varies across LPC, and that this pattern directly maps onto variations in functional connectivity using task-based and resting-state functional connectivity measures. Specifically, using ICA we demonstrated separable LPC functional connectivity networks, with each LPC subregion showing varying functional preference in a sentence, picture, and number sequence task. First, consistent with existing evidence LPC (Caspers *et al*. 2008; Uddin *et al*. 2010; Caspers *et al*. 2011; Mars *et al*. 2011; Cloutman *et al*. 2013), dorsal areas (dorsal PGa/IPS) demonstrated functional connectivity with the frontal executive network, whereas ventral LPC varied in an anterior-posterior direction with central ventral LPC (mid PGp) connecting with the DMN, anterior ventral LPC (ventral PGa) connecting with the fronto-temporal language network, and posterior ventral LPC (posterior PGp) connecting with the occipito-parietal visuospatial network. As PUCC would predict, these variations in functional connectivity were mirrored in terms of task activation profile. Dorsal LPC demonstrated a domain-general response, with equally strong positive activation for sentence, picture and number domains relative to rest. Whereas ventral areas varied along an anterior-posterior axis: specifically, the central AG (mid PGp), which functionally connected with the DMN, was equally deactivated by all three fMRI domains relative to rest; the anterior region that connected with the fronto-temporal language system showed positive activation only for the sentence task; and, the posterior region was part of the visual/SPL network and hence only responded to the picture sequences. Nevertheless, whilst this result is consistent with the predictions of the PUCC model, functional connectivity does not necessarily reflect the true underlying structural connectivity. Thus, one of the key aims of the current study was to examine the extent to which variations in activation patterns could arise from underlying structural variations white-matter connectivity.

### Technical issues/recommendations for LPC research

Previous studies have demonstrated that, in order to explore LPC functions and reveal interpretable findings, it is necessary to take certain factors into account within the design and analysis of any study:

#### 1) The direction of activation relative to rest

given the involvement of LPC in the DMN, it is of critical importance to consider whether a task positively or negatively engages the LPC relative to rest. Whilst many tasks generate deactivation in the AG, this is not always the case and the handful of activities that do positively engage the AG might be crucial sources of evidence about its true contribution (Humphreys and Lambon Ralph 2015). Contrasts between a cognitive task of interest vs. an active control condition are ambiguous because the difference could result from 1) greater positive activation for the task or 2) greater *deactivation* for the control. This issue becomes even more important when considering the impact of task difficulty on activation and deactivation in this region (see next). A straightforward expectation applied to almost all brain regions is that if a task critically requires the LPC then the LPC should be strongly engaged by that task. Indeed, this is the pattern observed in the anterior temporal lobe (ATL) where semantic tasks are known to positively engage the ATL relative to rest, whereas non-semantic tasks do not modulate/deactivate ATL (Humphreys *et al*. 2015). Perhaps one of the major motivations for considering task (de)activation relative to ‘rest’ is that ‘rest’ can be used as a common constant reference point across tasks. This is particularly important when conducting cross-domain comparisons. For instance, when one is examining a single cognitive domain it is possible to use a domain-specific baseline, i.e., a contrast a task that places strong demands on the particular cognitive system vs. a task with lower demands (e.g. remember > know in an episodic memory tasks, or words > non-words in a semantic memory task). Since the same is not possible across cognitive domains, rest acts a common constant for cross-domain comparisons, even if the true cognitive interpretation of ‘rest’ is unclear.

#### 2) Task-difficulty

Task-difficulty is important in two different ways. First, task difficulty correlates positively with activation in dLPC (dorsal AG/IPS) but negatively with the level of activation within vLPC or, put in a different way, the level of deactivation in vLPC (mid-AG) is positively related to task difficulty. Indeed, the dLPC and vLPC have often been shown to be anticorrelated in resting state data (Fox et al. 2009; Chai et al. 2012; Keller et al. 2013; Humphreys and Lambon Ralph 2017). Secondly, task-difficulty deactivations need to be accounted for when interpreting differences in ventral LPC areas. A ‘positive’ difference can be obtained in the AG simply by comparing easy > hard task conditions even for tasks that are entirely non-semantic, non-linguistic and non-episodic (Humphreys and Lambon Ralph 2017). One major limitation in the evidence for the semantic hypothesis is that apparent semantic fMRI effects could be explained by a difficulty confound (e.g., word > nonword, concrete > abstract). Indeed, it is known that the level of de-activation correlates with task-difficulty (Harrison et al. 2011; Gilbert et al. 2012; Humphreys and Lambon Ralph 2017), and it has been shown that one can both eliminate the difference between semantic and non-semantic tasks when task difficulty is controlled (Humphreys and Lambon Ralph 2017) and, more compellingly, also flip the typical ‘semantic’ effects (e.g. words > nonwords, concrete > abstract) by reversing the difficulty of the tasks or stimuli (Pexman et al. 2007; Graves et al. 2017).

#### 3) The importance of within-study comparisons

Reviews and formal meta-analyses of existing fMRI data have clearly identified overlapping areas of activations within the LPC (Humphreys and Lambon Ralph 2015). Whilst highly suggestive of domain-general computations in these regions, one needs within-participant comparisons to test these hypotheses further. Without such evidence, two alternative interpretations are possible: 1) True overlap across tasks implicating the region in a common neurocomputation (Corbetta and Shulman 2002; Walsh 2003; Cabeza *et al*. 2012;

Fedorenko *et al*. 2013; Humphreys and Lambon Ralph 2015, 2017) or 2) small-scale variability in function across the LPC which is blurred by cross-study comparisons or meta-analyses (Dehaene et al. 2003).

### The current study

The current study had two goals: 1) to compare alternative theories of LPC function directly within the same group of participants in an fMRI study, and 2) to examine the extent to which variations in the emergent task-activation patterns could arise from the underlying functional and structural connectivity across the LPC, as predicted by the PUCC model.

The first goal was addressed in Experiment 1: following some of the dominant proposals about LPC function (reviewed above), in an fMRI study we manipulated internally vs. externally directed attention, and episodic retrieval vs. semantic retrieval. Importantly, we considered the direction of activation/deactivation vs. rest as well as the extent to which the results can be explained in terms of variations in task-difficulty. There were four conditions, two involving internally-directed attention (semantic or episodic retrieval) and two involving externally-directed visual attention (real-world object decision or scrambled pattern decision). In each task the participant was presented with word-triads including a target word in the centre of the screen and two words below. In the semantic task, the participants indicated which item was semantically related to the target (e.g. moose: antlers or feathers). In the episodic task, the participants selected the feature that best matched the target items. The target items were verbal labels that corresponded to colour photographs that were viewed prior to the scan (e.g. bucket: green or red). Vividness ratings were made after each trial. In the object-decision tasks participants responded to a question about a picture presented on the screen (chair: blue red), and in the control task they indicated the direction of a scrambled picture that was shifted to the left or right side of the screen (see Figure 1). LPC activation was also contrasted with the activation in a known semantic region within the ATL.

**Figure 1:**
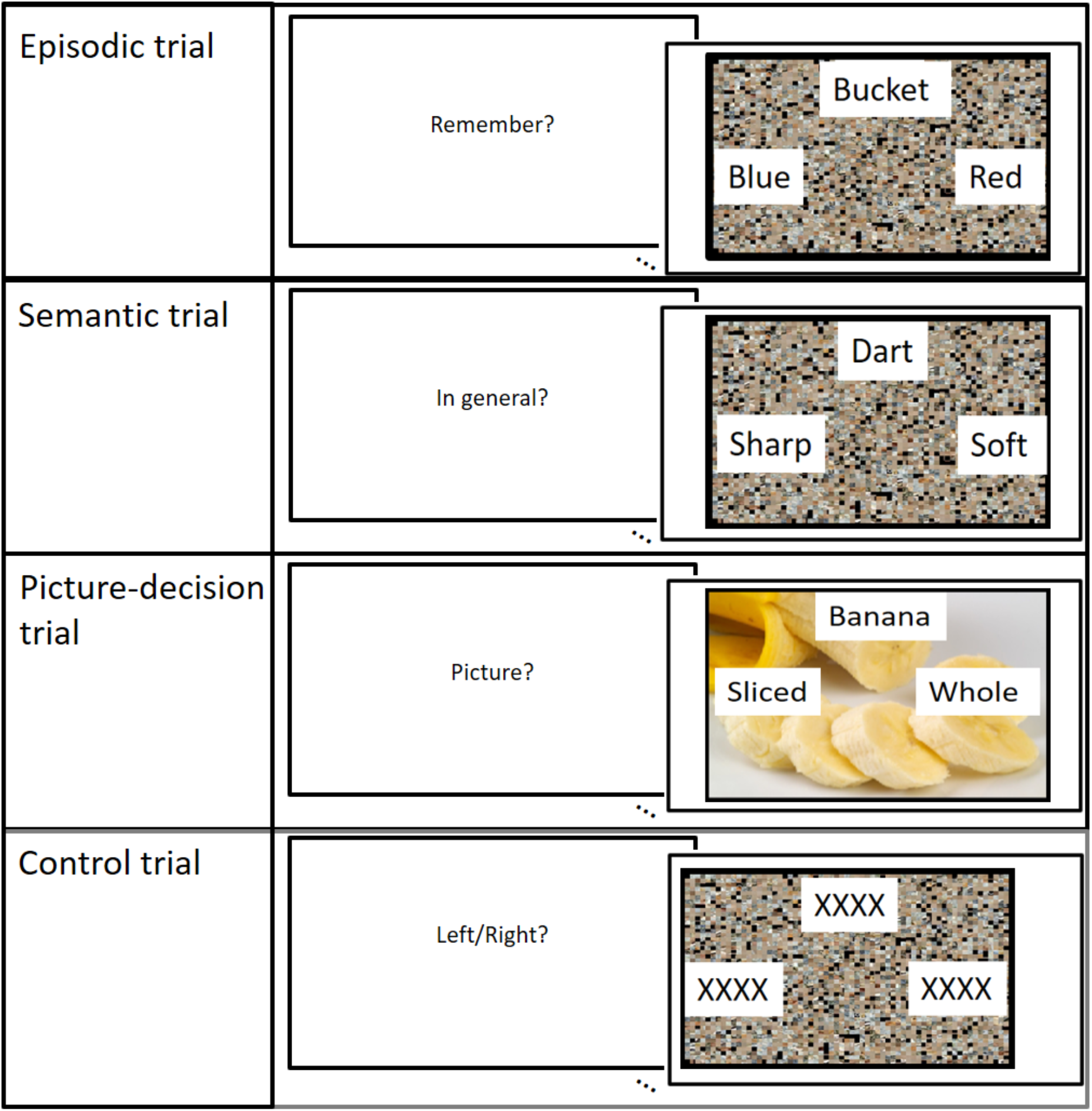
One example trial from each experimental condition

The second goal was addressed in Experiment 2: here we first sought to determine the correspondence between structural and functional connectivity measures of LPC. We have previously shown varying functional connectivity patterns across LPC subregions in the dorsal-ventral and anterior-posterior direction. In the current study, we directly compared the same functional connectivity pattern with structural connectivity from the same ROIs using DTI white-matter connectivity measures. Secondly, after establishing functional and structural input, we determined extent to which varying functional and structural input directly maps onto variations in the pattern of task-based activation in the same ROIs across two independent fMRI datasets.

## Methods

### Experiment 1: fMRI study

#### Participants

Twenty-two participants took part in the fMRI study (average age = 23.81, SD = 4.54; N female = 15). All participants were native English speakers with no history of neurological or psychiatric disorders and normal or corrected-to-normal vision.

#### Task design and procedures

There were four experimental tasks: episodic, semantic, picture-decision, and control. In each task the participant was presented with word-triads including a target word in the centre of the screen and two words below, one on the left and one on the right. The participants had to select the correct option by button-press. The words were presented on top of a scrambled or unscrambled picture depending on the condition. The trials lasted 4 seconds and were preceded by a 1.5 second instruction indicating the upcoming task. The trials were presented using an event-related design with the most efficient ordering of events determined using Optseq (http://www.freesurfer.net/optseq). Null time was intermixed between trials and varied between 0 and 26 seconds (average = 2.80 seconds, SD = 3.13) during which a fixation cross was presented. In total, 54 items were presented for each condition. The experiment was split into three runs (18 trials per condition), each run lasting 620 seconds, the order of which was counterbalanced across participants. An example trial from each task can be seen in Figure1.

#### Semantic task

Here the participants were presented with a target word (e.g., knife) and two-alternative possible features of the object (e.g., sharp vs. bendy), such as its typical function, colour, texture, shape etc. The participants were instructed to determine which alternative was correct. The words were presented on top of a scrambled picture.

#### Picture-decision task

Here the word triads were presented on top of a colour photograph of an object (e.g., a chair). The target word referred to a property of the picture (e.g., colour) and two alternative choices (e.g., blue and red). The participants were instructed to select the option that best matched the target feature of the object.

#### Episodic task

Immediately prior to the scan the participants were exposed to a selection of 54 colour photographs of objects (e.g., a bucket) and told that they would be required to remember aspects of the pictures during the experiment. Each photograph was presented for 10 seconds and the participants were asked to describe the picture in as much detail to ensure that the pictures were sufficiently encoded. In each trial of the experiment a target word would be presented (e.g., bucket) and the two alternative possible features of the remembered item (e.g., blue or red). The words were presented on top of a scrambled picture. The participants were instructed recall the feature that best described the target item. After a short jittered interval (varying from 0-1.5 seconds) the participants were given 3 seconds in which to rate the vividness of their memory of that particular item from 1-4 (1 = not vivid, 4 = very vivid). The episodic trial and the vividness rating were modelled separately in the general linear model.

#### Control task

In the control task the word-triads consisted of a string of Xs (e.g., xxxxxxxxx) on top of a scrambled picture. The picture was shifted slightly to the left or right. The participants had to indicate the direction of the shift. This acted as control for visual and motor activation.

#### fMRI acquisition parameters

Images were acquired using a 3T Philips Achieva scanner using a dual gradient-echo sequence, which has improved signal relative to conventional techniques, especially in areas associated with signal loss (Halai et al. 2014). 31 axial slices were collected using a TR = 2.8 seconds, TE = 12 and 35ms, flip angle = 95°, 80 × 79 matrix, with resolution 3 × 3 mm, slice thickness 4mm. B0 images were also acquired to correct for image distortion.

### fMRI data analysis

#### Preprocessing

The dual-echo images were first B0 corrected and then averaged. Data were analysed using SPM8. Images were motion-corrected and co-registered to the participants T1 structural image, and then spatially normalised into MNI space using DARTEL (Ashburner 2007). The functional images were then resampled to a 3 × 3 × 3mm voxel size and smoothed with an 8mm FWHM Gaussian kernel.

#### General Linear Modelling

The data were filtered using a high-pass filter with a cut-off of 190s and then analysed using a general linear model. At the individual subject level, each condition was modelled with a separate regressor and events were convolved with the canonical hemodynamic response function. Time and dispersion derivatives were added and motion parameters were entered into the model as covariates of no interest. At the individual level, each task condition was contrasted separately against rest, and entered at the second-level into separate one-sample t-tests to test for a significant group effects for each condition vs. rest, as well as one-way ANOVA to determine significant differences between tasks. A standard voxel height threshold p < .001, cluster corrected using FWE p < .05 was used for all group analyses.

#### Regressor analyses

It has been previously shown that LPC activity is modulated by task difficulty, with dorsal LPC showing a positive correlation with difficulty and ventral parietal cortex showing a negative correlation with difficulty (Harrison *et al*. 2011; Gilbert *et al*. 2012; Humphreys and Lambon Ralph 2017), this might be true on the task-level (harder tasks more strongly modulate activation) but also within each task at the item-level. In order to determine whether there were significant correlations with trial-to-trial difficulty, we added trial-wise RT as a parametric modulator to the GLM, 1) across all tasks, as well as 2) for each task independently. In addition to task difficulty, in terms of episodic memory retrieval, there is evidence that ventral LPC activation positively correlates with the self-reported vividness of the retrieved memory. In order to determine whether there were significant correlations with episodic vividness ratings, the item-level vividness ratings were added as a parametric modulator in a separate GLM.

#### Region of interest (ROI) analyses

The specific hypothesis regarding the AG and IPS were tested using ROIs taken from a previous large-scale multi-domain meta-analysis (AG and IPS) (Humphreys and Lambon Ralph 2015), and are therefore representative of the regions key regions highlighted in the literature. Note, the AG ROI corresponded to a 8mm sphere centred on the coordinates showing maximum likelihood of activation for both semantic and episodic meta-analyses, and the region showing domain-general executive processing in the IPS (Humphreys and Lambon Ralph 2015). In addition, one primary aim of the current study is to test the theory that the AG functions as multi-modal store of semantic information in a similar manner to the anterior temporal lobe (ATL). If true, the AG should show a similar pattern of task-related activation to that observed in the ATL. To test this, we compared task-activation in the AG vs. ATL ROIs. The ATL ROI was defined based on the results from a large-scale study that compared a variety of semantic tasks relative to non-semantic control tasks (Humphreys *et al*. 2015).

### Experiment 2

#### Functional connectivity analysis

The second key aim of the current study was to determine whether the pattern of functional connectivity found across IPL regions in a previous study is mirrored in the structural connectivity. The full details of the previous study can be found in Humphreys et al., (2020). Briefly, twenty-four participants completed a sequence processing task across three different domains: sentences, pictures, and numbers. On a given trial, a sequence of items (words, pictures, or numbers) was visually presented one item at a time and the participants indicated whether or not the sequence followed a coherent or incoherent structure. The pre-processed fMRI data were analysed in a group spatial ICA using the GIFT toolbox (http://mialab.mrn.org/software/gift) (Calhoun et al. 2001) to decompose the data into its components, and identify those that included the LPC. Four LPC components were identified, and functional labels were assigned to each based on the overlap with the built-in GIFT functional network template. The four components were labelled as follows: a DMN component, a fronto-parietal executive control component, a language component, and a visual-parietal component (see Figure 4 and description of components in results). The fronto-parietal executive component had a peak in dorsal LPC, within dorsal AG/IPS, whereas the other three components were located in ventral LPC. Specifically, the peak LPC coordinate for the DMN component was located in mid-AG (PGp), whereas the language component was slightly anterior (ventral PGa), and visual-parietal component was more posterior in posterior AG (posterior PGp). The same ICA networks were identified an independent resting-state ICA analysis suggesting the results are reliable.

#### Distortion-corrected diffusion weighted imaging and probabilistic fibre tracking

Diffusion-weighted images were acquired in 24 healthy volunteers (11 females; mean age 25.9, range 19–47) without any record of neurological or psychiatric disorders. This dataset has described previously and utilized for various tractography-related explorations (Binney et al. 2012; Cloutman et al. 2012; Bajada et al. 2016; Bajada et al. 2017; Jung et al. 2017; Jung et al. 2018). All participants were right-handed, as assessed by the Edinburgh Handedness Inventory (Oldfield 1971). Participants gave written informed consent to the study protocol, which had been approved by the local ethics committee.

A 3T Philips Achieva scanner (Philips Medical System, Best Netherlands) was used for acquiring imaging data with an eight-channel SENSE head coil. Diffusion weighted imaging was performed using a pulsed gradient spin echo-planar sequence, with TE = 59 ms, TR ≈ 11,884 ms, G = 62 mTm^−1^, half scan factor = 0.679, 112 × 112 image matrix reconstructed to 128 × 128 using zero padding, reconstructed resolution 1.875 × 1.875 mm, slice thickness 2.1 mm, 60 contiguous slices, 61 non-collinear diffusion sensitization directions at b = 1200 smm^−2^ (∆ = 29.8 ms, δ = 13.1 ms), 1 at b = 0, SENSE acceleration factor = 2.5. Acquisitions were cardiac gated using a peripheral pulse unit positioned over the participants’ index finger or an electrocardiograph. For each gradient direction, two separate volumes were obtained with opposite polarity *k*-space traversal with phase encoding in the left–right/right–left direction to be used in the signal distortion correction procedure (Embleton et al. 2010). A co-localized T2 weighted turbo spin echo scan was acquired with in-plane resolution of 0.94 × 0.94 mm and slice thickness 2.1 mm, as a structural reference scan to provide a qualitative indication of distortion correction accuracy. A high-resolution T1-weighted 3D turbo field echo inversion recovery image (TR ≈ 2000 ms, TE = 3.9 ms, TI = 1150 ms, flip angle 8°, 256 × 205 matrix reconstructed to 256 × 256, reconstructed resolution 0.938 × 0.938 mm, slice thickness 0.9 mm, 160 slices, SENSE factor = 2.5), was obtained for the purpose of high-precision anatomical localization of seed regions for tracking.

In order to directly compare the results from the DTI to those from the functional connectivity analysis, four regions of interest (ROIs) were identified for tractography based on the peak LPC coordinates from the four functional networks identified in the functional connectivity analysis: dorsal LPC (dorsal PGa/IPS), anterior-vLPC (PGa), mid-vLPC (PGp), and posterior-vLPC (PGp) (Humphreys, Jackson, *et al*. 2020). Note that the mid-LPC corresponds most closely to the DMN area, although the entire AG is often implicated. ROIs were defined as an 8mm radius sphere in the left hemisphere and transformed into each individual’s native diffusion space using the diffeomorphic anatomical registration through an exponentiated lie algebra (DARTEL) toolbox (Ashburner 2007) based on each participant’s T1-weigthed images.

DTI analysis was performed using unconstrained probabilistic tractography using the PICo software package (Parker et al. 2003), sampling the orientation of probability density functions (PDFs) generated constrained spherical deconvolution (CSD) (Tournier et al. 2008) and model-based residual bootstrapping (Haroon et al. 2009; Jeurissen et al. 2011). 20,000 Monte Carlo streamlines were initiated from each voxel within an ROI. Step size was set to 0.5 mm. Stopping criteria for the streamlines were set so that tracking terminated if pathway curvature over a voxel was greater than 180°, or the streamline reached a physical path limit of 500 mm.

The tracking results for each participants were spatially normalized into MNI space using the DARTEL toolbox. The brain regions associated with each fibre pathway were determined using brain masks from the AAL atlas or the Juelich histological atlas, which has a finer demarcation for the parietal cortex. The superior, middle, and inferior temporal lobe were further subdivided into anterior, middle, and posterior regions based on those used previously (Jung *et al*. 2017) since these regions have been shown to have large variations in connectivity profiles (Bajada *et al*. 2017; Jung *et al*. 2017). For each ROI, the atlas masks were overlaid over each participant’s tracking data and a maximum connectivity value (ranging from 0 to 20,000) between the ROI and each region of the brain was estimated. Thereby, we obtained a single probability estimate of a pathway between each pair of regions. These values were placed into an individual-specific matrix. Then, the connectivity matrices were subjected to a double threshold to ensure that only connections with high probability in the majority of participants were considered. For the first-level individual threshold, following the approach described by Cloutman et al (Cloutman *et al*. 2012), the λ-value of the Poisson distribution identified was used to determine a threshold value at p = 0.025. For the second-level group threshold, we used both a stringent (over 70% of participants, i.e., at least 17/24 participants) and a more relaxed (over 50% of participants, i.e., at least 12/24 participants) criteria for consistency.

#### fMRI ROI analysis

In order to determine the extent to which the results from the functional and structural connectivity analysis relate to functional differences in task-based activation, the same four LPC subregions (dorsal LPC (dorsal PGa/IPS), anterior-vLPC (PGa), mid-vLPC (PGp), and posterior-vLPC (PGp) were used as ROIs to examine the pattern of task-based activation from two fMRI studies 1) the sentence, picture, and number sequence task described above and reported in detail Humphreys et al. (2020), and 2) the current fMRI study reported in Experiment 1.

## Results

### Experiment 1

#### Behavioural results

On average the participants produced relatively few incorrect responses (average proportion incorrect (SD): episodic task = 0.13 (.05), semantic task = 0.06 (0.4), picture task = 0.09 (.03), and control task = 0.01 (0.1)) (see Figure 3). The average reaction time for each task were as follows (SD): episodic task = 2149ms (190), semantic task = 1888ms (182), picture task = 1842ms (231), and control task = 695ms (122). A one-way within-subjects ANOVA revealed that the tasks varied significantly in difficulty both in terms of accuracy (F(21) = 111.88, p < .001), and reaction time (F(21) = 1202.20, p < .001). Pairwise comparisons showed that the episodic task was significantly more difficult than all other tasks in terms of accuracy (all *t*s > 3, all *p*s < .006), the picture task was harder than the semantic task and control task (all *t*s > 3, all *p*s < .005), and the control task was easier than all other tasks (*t*s > 3, all *p*s < .001). The reaction time data mirrored the accuracy data (all *t*s > 6.58 and *p*s < 0001), with the exception that the semantic task and picture task did not differ (*t* = 1.28, *p* = .2). Given the known effects of task difficulty on IPS activation and AG deactivation, then – all other things being equal – one might have expected the greatest IPS activation, and the greatest AG deactivation for the episodic task followed by the picture task, then semantic and final the control task. Of theoretical relevance, whilst this was true in the IPS, this pattern was not the observed in AG.

#### fMRI results

Whole-brain analyses were performed using a standard voxel height threshold p < .001, cluster corrected using FWE p < .05. The episodic task > control task was found to activate bilateral lateral and medial frontal areas, dorsal (IPS) and ventral LPC (AG), medial parietal (precuneus), and posterior and medial occipito-temporal areas (see Figure 2). As with many other tasks with written word stimuli (Rice, Hoffman, et al. 2015; Rice, Lambon Ralph, et al. 2015) the semantic task > control task activation was mainly left-sided but overlapped with the episodic task in the lateral and medial frontal cortex, and posterior occipito-temporal areas although the activation for the semantic task spread more anteriorly along fusiform gyrus into the anterior temporal lobe, in congruence with its established role in semantic processing (Lambon Ralph *et al*. 2017). In contrast to the episodic task, no parietal activation was found for the semantic task (in fact the vLPC was more strongly activated by the control task relative to semantics) (Figure 2). Indeed, direct task comparisons showed these network differences to be statistically significant (Figure 2). Importantly, given the importance of the direction of activation relative to rest, the same AG region that is *positively* activated in the episodic task (episodic task > rest), is *deactivated* in the semantic task (rest > semantic task) (Figure 2).

**Figure 2.**
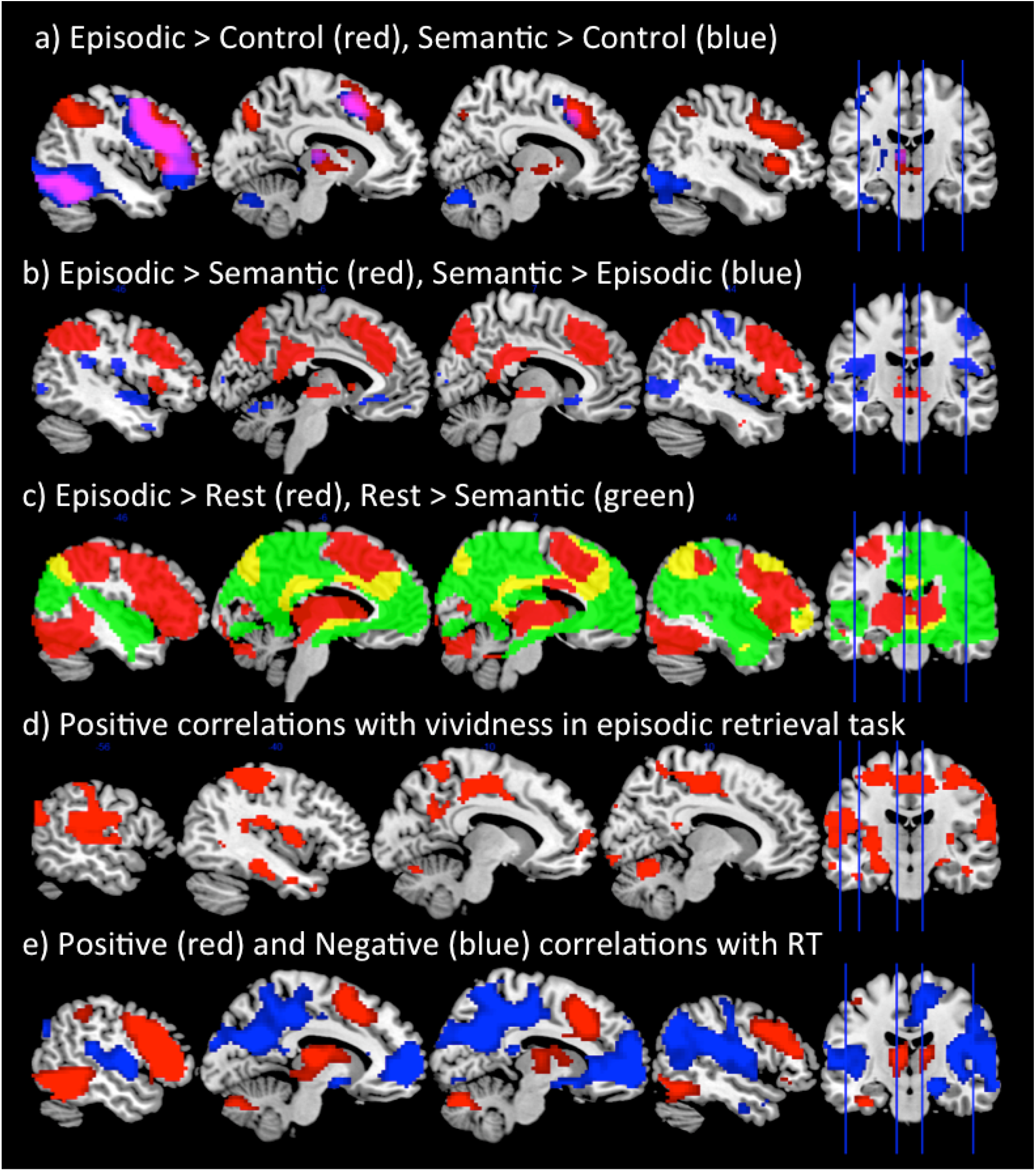
*A*: Whole-brain responses to the contrast of Episodic > Control (red) and Semantic > Control (blue). *B*: Whole-brain responses to the contrast of Episodic > Semantic (red) and Semantic > Episodic (blue). *C:* Whole-brain responses showing positive activation relative to rest for the episodic task (Episodic > Rest (red)), and deactivation relative to rest for the semantic task (Rest > Semantic (green)). This shows positive LTC activation for the episodic task but deactivation for the semantic task. *D*: The areas that positively correlated with vividness ratings during the episodic retrieval task, using a trial-wise parametric modulator. *E*: The network showing a positive correlation (red) and a negative correlation (blue) with task difficulty using a trial-wise parametric modulator. Analyses were performed using a standard voxel height threshold p < .001, cluster corrected using FWE p < .05.

Within AG, the results from the ROI analysis provided strong support that this region is primarily involved in episodic retrieval, rather than any other task. Despite being the hardest task, AG was found to be significantly *<u>positively</u>* activated relative to rest during episodic processing, as demonstrated by a one-sample t-test (t(21) = 2.30, p =. 03). In contrast, AG showed significant deactivation relative to rest during semantic retrieval (t(21) = -4.39, p < .00) thus strongly contradicting the semantic hypothesis. Indeed, semantic activation in the AG was found to also be lower than even the control task which required no semantic processing (t(21) = -2.84, p = .01) (Figure 3). This provides clear evidence that the AG is engaged by episodic retrieval, but is disengaged during semantic retrieval. Unlike the AG, however, the ATL (Figure 3), a known semantic region, showed a very different pattern of activation: strong positive activation relative to rest across the three experimental tasks (since all tasks required semantic processing) (all *t*s> 7.34, *p*s < .001), but no modulation for the control task (*t*=0.86, *p*=0.4). In further support of the role of AG in episodic retrieval, the network was found to be highly correlated with the vividness ratings. A whole-brain analysis showed a strong correlation with vividness within lateral and medial parietal cortex, as well as medial frontal areas (Figure 2). This network shows close correspondence to the DMN. Therefore, AG activation appears closely related to vivid episodic retrieval, rather than semantic retrieval.

**Figure 3.**
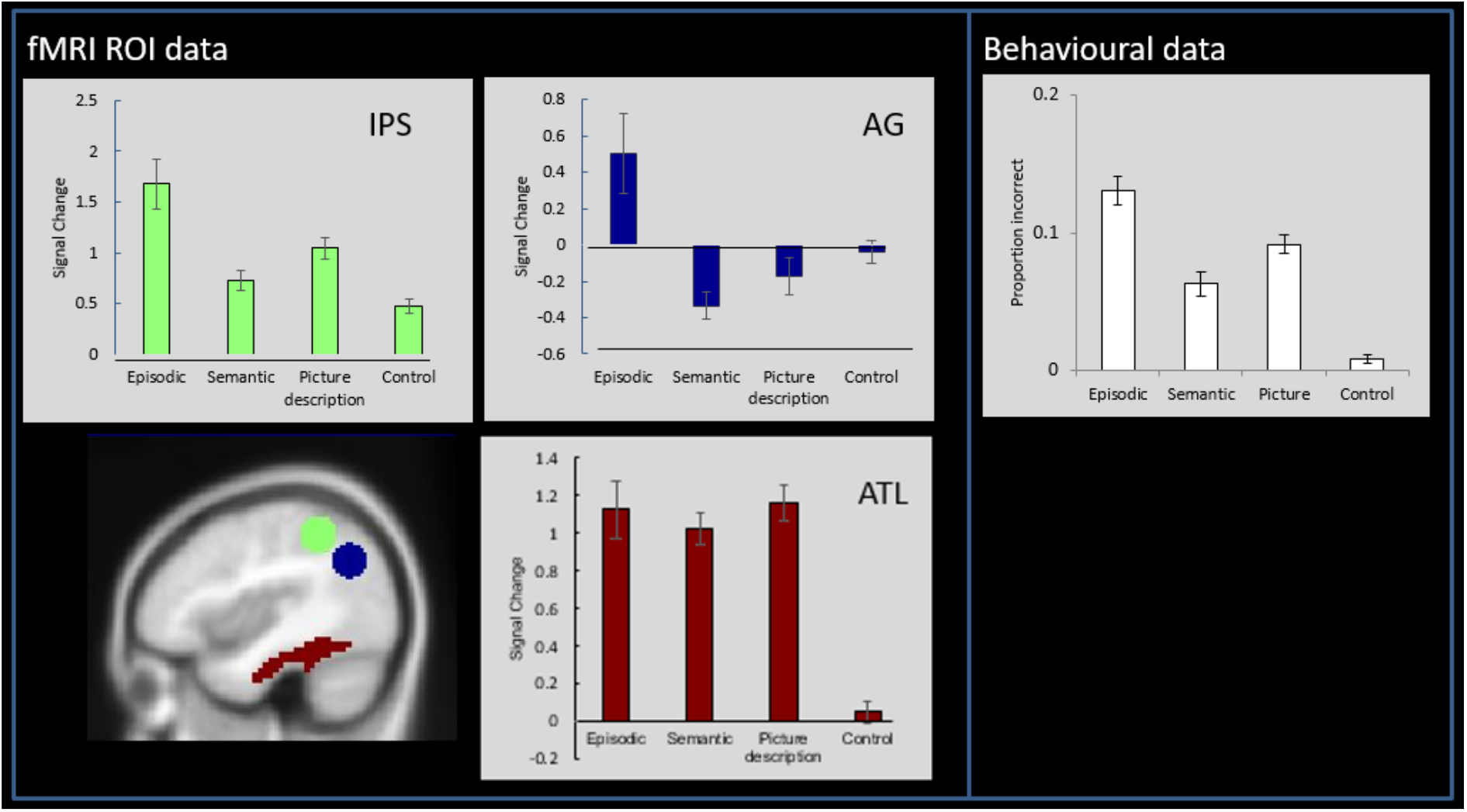
Left panel: ROI analyses showing mean activation relative to rest for each condition within the IPS, AG, and ATL. The AG and IPS ROIs were defined based on the results from a large-scale multidomain meta-analysis (Humphreys and Lambon Ralph 2015). The ATL ROI was defined based on the region identified by a study that contrasted the activation from a variety of semantic tasks relative to non-semantic control tasks (Humphreys *et al*. 2015). Right panel: The behavioural data showing the average proportion of incorrect responses during the fMRI task. Note. The behavioural data closely mirrors the fMRI activity in the IPS.

The IPS was found to be positively activated relative to rest for all tasks (all ts > 5.44, p < .000) (Figure 3). Indeed, activation closely mirrored task difficulty (Figure 3), with significantly greater activation for the episodic task relative to all others (all ts > 2.70, p < .01), the picture task showing greater activation compared to the semantic task and control task (all *t*s > 4.20, all *p*s < .001), and the control task showing the weakest activation compared to all tasks (*t*s > 2.19, all *p*s < .001).

In order to investigate the relationship between task difficulty and activation across experimental items, we added RT as a parametric modulator to GLM analysis to examine the positive and negative correlation with trial difficulty in a whole-brain analyses. This revealed that a large network of dorsal parietal cortex, as well as lateral frontal, and posterior temporal areas were strongly positively correlated with task difficulty. In contrast, ventral parietal cortex, as well as the wider DMN in medial frontal and parietal cortex were strongly negatively related to task difficulty thus supporting the notion that this network is disengaged when a task becomes harder to perform (Figure 2). Note that the exception to this rule is during episodic retrieval, which was the hardest task behaviourally but nevertheless most strongly engaged ventral parietal cortex. Indeed, when the same whole brain correlation analysis was performed but only including the episodic retrieval trials no correlation was found with task difficulty within the DMN (even at the very lenient threshold *p* < .05 uncorrected).

To summarise, the AG was actively engaged by the episodic retrieval task, and this positively correlated with the vividness of the memory. In contrast, the semantic task, picture-decision task, and control task all resulted in AG deactivation, and the level of deactivation was proportionate to task difficulty. The IPS, showed the reverse response, with increasing activation for increased task difficulty.

### Experiment 2

#### Functional connectivity results

Four LPC functional networks were identified from the functional ICA analysis (Figure 4). These included one dorsal LPC component which encompassed a fronto-parietal executive control network with peaks in dorsal AG/IPS, left lateral frontal, pMTG, and posterior superior frontal gyrus (referred to as the executive network from here on). The remaining three components implicated ventral LPC, with an anterior-posterior organisation. These included 1) A DMN component with a peak in mid-AG (PGp), precuneus (PCC), medial frontal, mid-middle temporal gyrus (MTG)), 2) a component that clearly resembled the language network including the anterior AG (PGa), left IFG, and large section of the temporal lobe (STG and MTG) (Vigneau et al. 2006), and 3) in posterior AG (posterior PGp), a visual-parietal network that involved visual cortex, SPL and PGp.

**Figure 4.**
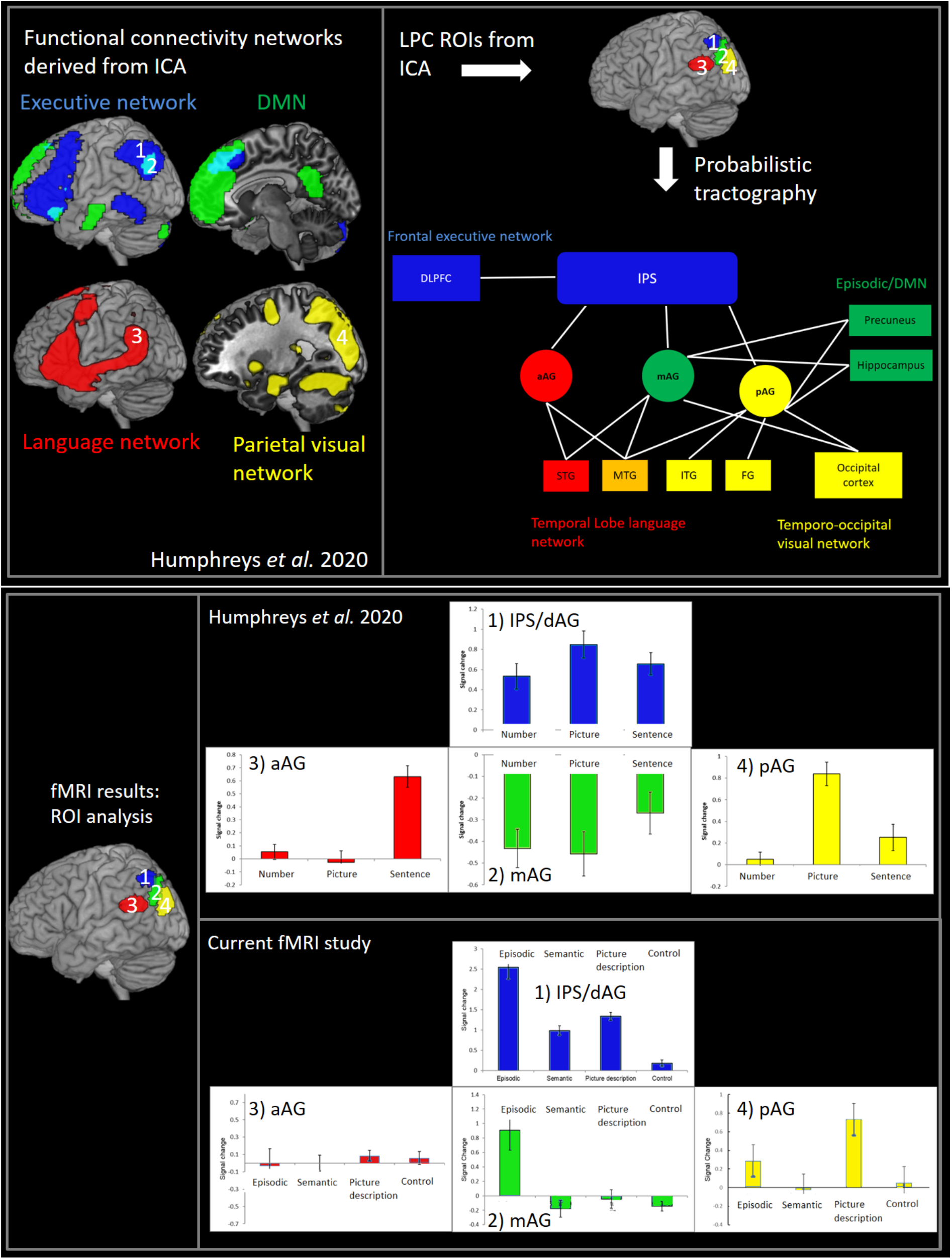
Top left: The four LPC functional connectivity networks derived from ICA. Top right: The results from the DTI analysis, using the ICA-derived LPC seed regions. Bottom panel: The results from two fMRI studies using the LPC ROIs.

#### DTI results

In order to examine the extent to which the functional connectivity networks reflect direct structural connections, the coordinates for the four peak LPC sub-regions (dorsal AG/IPS, mid-AG, anterior-AG, and posterior-AG) from the functional ICA were used as ROIs for the DTI analysis. Note that the mid-AG and dorsal AG/IPS ROIs overlap with the AG and IPS ROIs used in Experiment 1, respectively.

##### Dorsal connectivity

Consistent with the role of dLPC in executive processing the IPS/AG ROI showed long-range connectivity with lateral frontal lobe, specifically, dorsolateral frontal cortex (DLPFC) and IFG (BA44) which are known to be involved in top-down executive control (Figure 4, Table 1).

**Table1.** Group-level connectivity matrix. Bold font indicates that the connection probability was over 70% for the group analysis. The individual threshold was set at 2.5%.

|  |  | Dorsal<br>IPS | Anterior | Ventral<br>Middle | Posterior |
| --- | --- | --- | --- | --- | --- |
| Temporal lobe | mMTG |  | **92** |  |  |
|  | pSTG |  | **96** |  |  |
|  | pMTG |  | **96** |  |  |
|  | pITG |  |  | 58 | 54 |
|  | Hippocampus |  |  | **83** | **92** |
| Frontal lobe | DLPFC | **71** | **71** |  |  |
|  | BA44 |  | 50 |  |  |
| Parietal lobe | S1 | **100** | **75** | **88** | 58 |
|  | 5Ci |  |  |  |  |
|  | 5M |  |  |  |  |
|  | 5L |  |  | 63 |  |
|  | 7PC | **100** | 63 | **92** | 67 |
|  | 7A | **96** |  | **96** | **79** |
|  | 7P |  |  | **83** | **71** |
|  | 7M |  |  | 54 | **75** |
|  | IPS1 | **100** | **75** | **92** |  |
|  | IPS2 | **100** | 67 |  |  |
|  | IPS3 | **100** |  | **96** | 63 |
|  | PFo | 63 |  |  |  |
|  | PFt | **83** |  |  |  |
|  | PF | **96** | **88** |  |  |
|  | PFm | **75** | **88** |  |  |
|  | PFcm | **96** | **88** |  |  |
|  | PGa | 67 | **92** | **88** |  |
|  | PGp |  | 63 | **100** | **83** |
|  | Precuneus |  |  | **88** | **100** |
| Occipital lobe | Superior OCC |  |  | **92** | **100** |
|  | Middle OCC | **79** | 67 | **100** | **100** |
|  | Inferior OCC |  |  | 50 | **83** |

##### Anterior-posterior gradient within the ventral parietal lobes

Within the vLPC, there was a graded variation in connectivity between anterior and posterior parietal cortex control (Figure 4, Table 1). The anterior vLPC showed connectivity with the temporal lobe areas implicated in language processing (Vigneau *et al*. 2006; Binder *et al*. 2009; Lambon Ralph *et al*. 2017). In contrast, moving in a posterior direction, in mid vLPC the connectivity tended to become stronger with areas associated with the DMN and episodic memory, including the inferior temporal cortex and the hippocampus, as well as large portions of the precuneus (Sestieri et al. 2011; Rugg and Vilberg 2013). Finally, the posterior vLPC connected with areas including the medial parietal cortex and occipital lobe associated with visual processing and spatial attention (Corbetta and Shulman 2002; Zacks 2008). Together these results appear highly consistent with the functional networks identified in the functional ICA.

### fMRI ROI analyses

A core assumption of the PUCC model is that differences in the underlying structural connectivity inputs across LPC give rise to variations in the emergent functional activation profile across the region. To test this assumption we examined the functional activation within each of the four LPC ROIs in two independent fMRI datasets: 1) the sentence, picture, and number sequence task, reported previously (Humphreys et al., 2020), and 2) the fMRI data from Experiment 1 involving the episodic, semantic, picture-description, and control task (see Figure 4). For the first fMRI dataset, dorsal AG/IPS and mid-AG ROIs exhibited opposing directions of activation relative to fixation; activation for the dorsal AG/IPS, which part of the executive network, was significantly greater than rest for all conditions (one-sampled t-test, all ts > 3.49, ps < .002). In contrast, the ventral mid-AG, which is part of the DMN/episodic memory network, showed significant negative activation for all conditions (one-sampled t-test, all ts > -3.68, ps < .002). The anterior and posterior ventral AG showed a different pattern: the anterior ventral AG, which formed part of the language network, was only activated for the sentence task, showing significantly positive activation for the sentence conditions only (ts > 4.72, ps < .001, d = 1.01), with the picture and number conditions showing no difference from zero (ts < .92, ps > .37). In contrast, the posterior ventral AG, which formed part of the visual-parietal network was specifically engaged by the picture task only (ts > 6.35, ps < .001, d = 1.42), with no modulation of the sentence and number tasks (ts < 2, ps > .05).

For the second fMRI dataset from the current study (Figure 4), like in the first, the dorsal AG/IPS was found to be positively activated by all tasks relative to rest (all ts > 8.50, p < .001). In contrast, whereas the sentence, picture, and number sequences deactivated the mid-AG in the first dataset, here mid-AG was positively activated for the episodic task (t(21) = 3.28, p =. 005) (all other tasks showed no difference from zero (all ts < -1.55, ps >. 14). The fact that the mid-AG is positively engaged by the episodic retrieval task in isolation is consistent with what one would expect if it functions as part of the DMN/episodic retrieval network, since no other task required the retrieval of information from episodic memory. The posterior AG, which was positively engaged by picture sequences in the first fMRI dataset, similarly showed positive activation relative to rest for the picture-description task (t(21) = 4.70, p <. 001) but no other (all ts < 1.38, ps >. 18), thereby supporting the role of this region as part of the visual-parietal network. Finally, whereas the anterior AG was engaged by the sentence sequence task in the first fMRI dataset, no modulation was found for any task in the second dataset which only included single-word items (all ts < 1.40, ps >. 18), this might thereby imply a greater role of this region in sentence/multi-item rather than single-word processing.

## Discussion

<u>Summary of main results: The results from Experiment 1 showed that the</u> dorsal and ventral LPC were positively engaged during episodic retrieval (but not in any other task), and this activation was found to correlate positively with the vividness of the episodic memory. The vLPC (AG) showed strong deactivation compared to rest during semantic memory retrieval (indeed the LPC was less engaged by the semantic task than the control task that involved little/no semantic processing). This provides compelling evidence against theories that posit a role in semantic processing, or in all forms of internally-directed thought. This pattern contrasts sharply with the results from the ATL, which showed strong positive activation for the semantic, episodic, and the picture-description task, since all three tasks necessitate the retrieval of information from semantic, but no modulation for the control-task (which had no semantic memory requirements). Indeed, if the AG served a primarily semantic function, one would predict a similar pattern of task-related activation to the ATL. In terms of task difficulty, with the exception of the episodic task, activation within the vLPC (AG), as part of the wider DMN, showed a strong negative correlation with reaction time thereby suggesting that this region is “turned-off” when a task becomes increasingly difficult. This is in contrast to the dLPC, as part of a wider multi-demand fronto-parietal network, which increased activation in relation to task-difficulty. Critically, the episodic retrieval task was the only task to activate both the dLPC and vLPC regions concurrently. These data were complemented by the results from Experiment 2, which showed that in terms of functional and structural connectivity, dLPC forms part of a fronto-parietal network, and was positively engaged by all fMRI tasks. The vLPC showed an anterior-posterior variation in connectivity. Specifically, the mid vLPC region (corresponding to the mid-PGp subregion of AG associated with the DMN) connected with areas associated with episodic memory retrieval (hippocampus and precuneus), and was only actively engaged during the episodic-retrieval task consistent with its role as part of the episodic retrieval network (Sestieri *et al*. 2011; Rugg and Vilberg 2013). In contrast, the anterior vLPC (PGa) showed functional and structural connectivity with temporal lobe language processing areas (Vigneau *et al*. 2006; Binder *et al*. 2009; Lambon Ralph *et al*. 2017), and was found to respond only to a sentence processing task, consistent with existing evidence from sentence processing studies (Humphreys and Lambon Ralph 2015; Branzi et al. 2020; Humphreys, Jackson, *et al*. 2020). Whereas, the posterior vLPC (posterior PGp) connected with the occipital lobe and medial parietal areas associated with visual attention (Corbetta and Shulman 2002; Zacks 2008), and was only actively engaged by the picture-sequence and picture-description task. Together these results fit with the PUCC model that suggests a shift in the functional engagement of vLPC based on varaitions in the underlying structurally connectivity of the network (see Figure 4 for a schematic model).

### Implications for semantic theories of LPC function

The current data provide clear evidence that the AG is not engaged during semantic retrieval. How can this result be aligned with evidence showing AG apparent sensitivity to semantic contrasts (Binder *et al*. 2009)? One key explanation is based on the type of contrast being performed: typical contrasts being words > nonword, or concrete > abstract. A meta-analysis of semantic vs. non-semantic studies that used other forms of contrast, found no evidence of AG engagement despite engagement of the wider fronto-temporal semantic network (Visser et al. 2010), and another study found stronger AG engagement during non-verbal tongue movements compared to meaningful-speech (Geranmayeh et al. 2012). As highlighted in the Introduction, when interpreting these data it is important to take into account two variables: 1) the polarity of activation relative to rest and 2) difficulty-related differences.

With regard the first point, the semantic retrieval task showed deactivation relative to rest. It is of course difficult to interpret “rest”, as it could involve spontaneous semantic and linguistic processing (Binder et al. 1999) (although one could also make the argument that the same is also true for episodic retrieval (Buckner *et al*. 2008; Andrews-Hanna 2012) yet episodic memory tasks positively engage the AG, as demonstrated here and elsewhere (Humphreys and Lambon Ralph 2015)). Importantly, whilst AG deactivates for semantic and non-semantic tasks, other key semantic areas do not show the same pattern as AG; for instance, the anterior temporal lobe is positively engaged during semantic tasks but deactivated by non-semantic tasks (Humphreys *et al*. 2015; Humphreys and Lambon Ralph 2017).

With regard the second point, AG deactivation is known to relate to task difficulty (Hahn et al. 2007; Mason et al. 2007; Humphreys and Lambon Ralph 2017), with increasing deactivation for harder tasks. Indeed, the contrast of word > nonword and concrete > abstract typically involve comparing an easier task vs. a harder task. Compellingly, the same word > nonword and concrete > abstract contrasts can be inverted by reversing the difficulty of the task/stimuli (Pexman *et al*. 2007; Graves *et al*. 2017). Indeed, when task difficulty is directly manipulated in a semantic and visuo-spatial task one observes a main effect of task difficulty (easy vs. hard) in the AG but no semantic vs. non-semantic difference, whereas the IPS shows the reverse pattern of difficulty-related activation (hard vs. easy) (Humphreys and Lambon Ralph 2017).

### Implications for episodic memory retrieval

The episodic retrieval task showed positive activation in both the AG and IPS despite being the most difficult task; making it the exception to the difficulty-related activation pattern. The influence of connectivity into various IPC areas, embraced in the PUCC framework (see below), can explain these episodic findings in the ventral and dorsal parietal cortex. Specifically, the mid-PGp region of the AG connects with the hippocampus and precuneus, operating as part of a wider episodic retrieval network. Based on evidence that the patients with AG damage are not profoundly amnesic, unlike those with damage to the medial temporal lobe (Berryhill et al. 2007; Simons et al. 2008; Humphreys, Lambon Ralph, *et al*. 2020), we propose that episodic information stored elsewhere in the system is temporally buffered online during episodic retrieval. Indeed the notion of an AG episodic buffer has been proposed elsewhere (Wagner *et al*. 2005; Vilberg and Rugg 2008). This notion is consistent with observation that patients with parietal damage lack clarity or vividness of episodic memories, as one might predict from a deficit in buffering multi-modal contextual information (Davidson et al. 2008; Shimamura 2011; Yazar et al. 2014; Moscovitch et al. 2016; Bonnici et al. 2018; St. Jacques 2019), as well as fMRI studies showing that the level of AG activation varies depending on the extent to which information is retrieved from episodic/autobiographical memory compared to semantic memory (Brown et al. 2018). In addition, due to its connectivity with DLPFC executive systems, the IPS takes on a domain-general ability for selection/manipulation of internally-buffered information – which is required in many episodic and semantic tasks. Indeed, previous studies of episodic retrieval have also suggested that the IPS plays an executive role in decision-making during episodic tasks (Gonzalez et al. 2015; Sestieri *et al*. 2017). Overall, we propose that episodic retrieval is an active construction process where the memory is reconstructed online thus needs both buffering of information from episodic-related areas and also executively-demanding shaping, thereby recruiting both dLPC and vLPC systems.

### A unifying account of lateral parietal cortex functions

By exploring both the variation in functional response and connectivity profiles across the LPC, the current study found evidence in favour of the neurocomputational principles embraced by the PUCC model (Humphreys and Lambon Ralph 2015; Humphreys, Jackson, *et al*. 2020; Humphreys, Lambon Ralph, *et al*. 2020). According to the PUCC model, the LPC does not support long-term stored information per se but rather is an online temporary buffer of multi-modal spatio-temporal input. Indeed, this hypothesis appears consistent with other proposals that AG acts a temporary buffer (Wagner *et al*. 2005; Vilberg and Rugg 2008), or a “schematic-convergence zone” which binds information, if we assume that this binding is temporary (Shimamura 2011; Wagner et al. 2015). An online buffer would seem to be a necessary neurocomputation for the construction of internal models of the world, reconstruction of autobiographical memories, or the envisioning of possible future events, and, perhaps, for the ongoing buffering of combinatorial meaning generated over a time-extended period (Hasson et al. 2008; Lerner et al. 2011; Ramanan and Bellana 2019). We would predict that if the LPC was operating as a “buffering-system” then the content of the system would align with what participants buffer at each temporal interval. Indeed, consistent results have been found from episodic memory literature using MVPA (Wagner *et al*. 2015; Lee and Kuhl 2016), whereby the episodic content of a person’s current recall (in this case the visual features of a face) directly align with decoding in the AG (Lee and Kuhl 2016).

A key notion of PUCC is that the expressed functions of a cortical region will reflect the combination of the local computation and its pattern of connectivity. This tenet has been formally demonstrated by computational models, whereby the resultant “behaviour” of a processing unit depends not only on its local computations, but also on its long-range connectivity (more recently referred to as ‘connectivity-constrained cognition – C3’: (Lambon Ralph et al. 2001; Plaut 2002; Chen et al. 2017)). How this might apply to the LPC is sketched out in Figure 5. In its most simple form (left panel), the LPC might have a single local computation – an online multi-modal buffer of internal and external sequentially-experienced information. Computational models with recurrent connections (shown as a generic ‘Elman’ network in Fig 4A) show this general property and the same type of model can be applied to any time-extended series of inputs and outputs (Elman 1990; Botvinick and Plaut 2006). Accordingly, across a larger region such as the LPC, even if the local buffering computation itself was the same, the emergent observed cognitive function will depend on what sources of information and influences arrive at each subregion: i.e., the differential connectivity pattern. Figure 5 (right panel) shows a schematic of the varying pattern of connectivity to LPC subregions. We can consider the influence of this connectivity profile in two steps. First, the long-range connectivity from executively-related DLPFC primarily terminates in the IPS/dorsal LPC region (Crowe et al. 2013) and not the ventral parietal cortex (VPC). This should generate a fundamental, counterpointed difference in the emergent observed functions: in receiving top-down signals from frontal executive processing areas, dLPC (IPS) will operate as a domain-general executive system engaged in the selection/manipulation processes on internally-buffered information (Fedorenko *et al*. 2013; Humphreys and Lambon Ralph 2015, 2017). The IPS region itself connects to all subregions of the VPC and thus can become involved in any particular activity irrespective of information type (c.f., the DLPFC and IPS are two key components of the ‘multi-demand’ system: (Fedorenko *et al*. 2013; Assem et al. 2020). In contrast to the IPS, most of the VPC does not seem to receive this same level of DLPFC connectivity and thus its underlying buffering will not be so executively penetrated; i.e., its buffering will be more “automatic” (e.g., a distinction envisaged in the differentiation between the central executive and ‘slave’ subsystems in classical models of short-term working memory: (Broadbent 1982; Baddeley 2000, 2003). More generally the differential connectivity of DLPFC to IPS and not VPC might also help to explain why in many situations dLPC and vLPC show anti-correlated functional activity (Fox *et al*. 2009; Chai *et al*. 2012; Keller *et al*. 2013; Humphreys and Lambon Ralph 2017); whereby the dLPC is increasingly activated and the vLPC increasingly deactivated based on task difficulty (Humphreys and Lambon Ralph 2017). When engaged in a goal-oriented task, on-going automatic information accumulation in VPC subregions would presumably be unhelpful/disruptive unless this input is necessary for task performance. In this case, activation in the irrelevant VPC subregions would be suppressed/deactivated (Humphreys, Jackson, *et al*. 2020). This would be especially needed with increasing demands on task performance, thereby explaining the common observed pattern of anticorrelated activation modulated by task difficulty (Humphreys and Lambon Ralph 2017).

**Figure 5.**
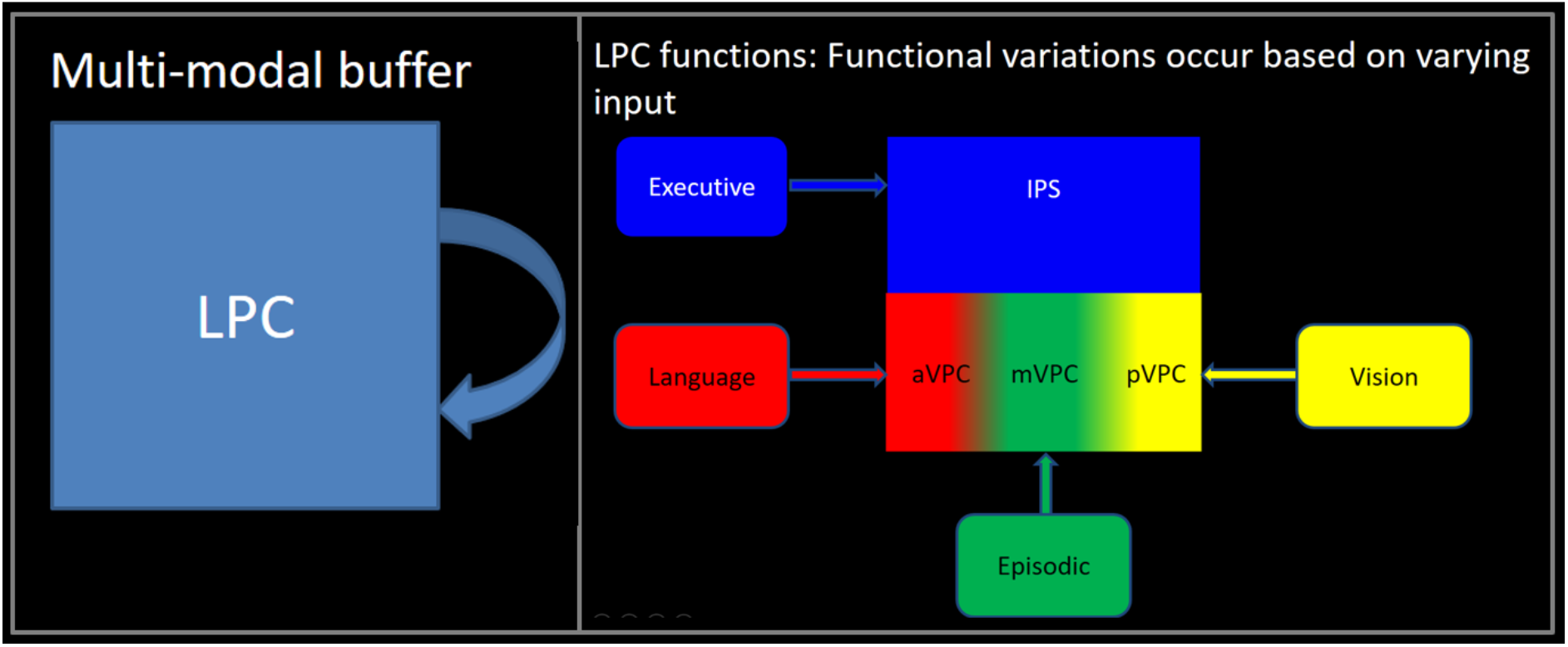
A schematic illustration of the assumptions of the PUCC model: 1) The LPC acts as a multi-modal buffer in terms of its basic underlying neurocomputation (left panel); and then 2) the varying long-range connectivity to LPC influences the emergent function of each subregion.

Different functions across the VPC itself should emerge given that there are differences in connectivity of VPC subregions to distinct neural networks (Fig. 4 and 5) (Nelson et al. 2012; Daselaar et al. 2013; Humphreys and Lambon Ralph 2015; Humphreys, Jackson, *et al*. 2020). Specifically, the more anterior VPC through its primary connections to pSTG and pMTG becomes involved in sound, phonological and language processing (Griffiths and Warren 2002; Humphreys and Lambon Ralph 2015). The most posterior VPC subregion is the most heavily influenced by the dorsal connectivity from visual regions, consistent with its involvement in visuospatial processing (Corbetta and Shulman 2002; Zacks 2008; Humphreys and Lambon Ralph 2015, 2017; Humphreys, Jackson, *et al*. 2020), whilst the mid-AG region’s involvement in episodic tasks seems entirely consistent with its connectivity through to key nodes of the extended episodic network (Sestieri *et al*. 2011; Rugg and Vilberg 2013).

To conclude, we propose a unified model of LPC function in LPC acts as multi-modal buffer of information. Despite a common core mechanism, graded subdivisions in how this function is expressed can arise based on varying long-range connectivity inputs to dorsal-ventral and anterior-posterior areas.

## Acknowledgements

This research was supported by an MRC Programme grant to MALR (MR/R023883/1) and an intramural award (MC_UU_00005/18).

